# Cell-based screen identifies human type I interferon-stimulated regulators of *Toxoplasma gondii* Infection

**DOI:** 10.1101/2022.01.02.474728

**Authors:** Anton Gossner, Anna Raper, Musa A. Hassan

**Affiliations:** Division of Infection and Immunity, The Roslin Institute, The University of Edinburgh, Edinburgh, United Kingdom

## Abstract

Macrophages activated with interferons (IFNs) respond with transcriptional changes that enhance clearance of intracellular pathogens such as *Toxoplasma*, a ubiquitous apicomplexan parasite that infects more than a billion people worldwide. Although IFNs generally inhibit *Toxoplasma*, the parasite can also induce components of the host IFN signalling pathway to enhance survival in host cells. Compared to the type II IFN gamma (IFNγ), the role of type I IFNs in macrophage response to *Toxoplasma* is relatively not well characterized. Here, using fluorescent *Toxoplasma* and a CRISPR/Cas9 knockout library that only targets interferon-stimulated genes (ISGs), we adapted a loss-of-function flow cytometry-based approach to systematically identify type I ISGs that control *Toxoplasma* growth in THP-1 cells, a human macrophage cell line. The system enabled the rapid screening of more than 1900 ISGs for type I (IFNα)-induced inhibitors and enhancers of *Toxoplasma* growth in THP-1 cells. We identified 26 genes that are associated with *Toxoplasma* growth arrest out of which we confirmed *MAX, SNX5, F2RL2*, and *SSB*, as potent IFNα-induced inhibitors of *Toxoplasma* in THP1 cells. These findings provide a genetic and experimental roadmap to elucidate type I IFN-induced cell-autonomous responses to *Toxoplasma*.

## INTRODUCTION

*Toxoplasma gondii*, a zoonotic protozoa that infects over a billion people worldwide, is one of the most common foodborne parasites with the greatest global impact^1^. *Toxoplasma* is a major cause of coma and death in HIV/AIDS patients as well as childhood blindness and defects in foetal brain development. Although infection in healthy individuals is mostly asymptomatic, latent parasites can reactivate in immunocompromised individuals and cause severe diseases such as encephalitis, hepatitis, and myocarditis^2^. While not yet proven, *Toxoplasma* is also causally linked with several neurological and behavioural disorders including schizophrenia^3^. Current anti-*Toxoplasma* drugs are not well tolerated and are ineffective during chronic infections, making the development of novel *Toxoplasma* control strategies an important priority for biomedical research.

Interferons (IFNs), a large family of related cytokines that form an integral part of the innate immune system – the first line of defence against invading pathogens, are known to play an important role in *Toxoplasma* pathogenesis. Activation of host cells, including macrophages by these cytokines leads to the upregulation of many effector molecules that can kill or inhibit *Toxoplasma*. They can also shape the adaptive immune response to *Toxoplasma* by triggering release of cytokines and chemokines. Developing a better understanding of the IFN response, and how *Toxoplasma* counteract their effects can, therefore, have important implications for how we treat *Toxoplasma* infections. Currently, three main types of IFNs are known: type I, II, and III^4^. IFN gamma (IFNγ) – the only type II IFN – induces the transcription of hundreds of interferon-stimulated genes (ISGs) by activating Janus Kinase 1 and 2 (JAK1/2) to phosphorylate homodimers of the signal transducer and activator of transcription 1 (STAT1). Type I IFNs (IFNα and IFNβ), induces the expression of ISGs with canonical IFN-sensitive response element (ISRE) by activating the phosphorylation of STAT1 and STAT2 heterodimers to interact with interferon regulatory factor 9 (IRF9)^5^. IFNγ is a well-characterized mediator of cell-autonomous immunity against *Toxoplasma* and other intracellular pathogens in vertebrates^6–8^. IFNγ can also inhibit or initiate the development of *Toxoplasma* cysts in the central nervous system and muscle tissues. For example, IFNγ induces GTPases such as p47 immunity-related GTPases (IRGs) and guanylate binding proteins (GBPs) that destroy the parasitophorous vacuole membrane (PVM) leading to the death of the parasite within^9^, or nitric oxide (NO) that inhibit parasite replication and initiate the developmental switch from tachyzoites to bradyzoites^10^. Mice lacking IRGs, GBPs, or NO are highly susceptible to *Toxoplasma*^11^. Expectedly, *Toxoplasma* virulence factors target components of the IFN signalling pathway. Although important insights into host-*Toxoplasma* interactions have been gained from studies in mice, there are several differences between murine and human immune systems. For example, humans lack functional toll-like receptors (TLRs) 11 and 12^12^, and human macrophages do not produce NO in response to IFNγ or *Toxoplasma*^13^, which are critical for the detection and inhibition of *Toxoplasma* in mice. Nevertheless, IFNγ plays a central role in human cell-autonomous responses against *Toxoplasma*. For example, IFNγ-induced tryptophan degradation by indole-2,3-dioxygenase (*IDO*) expression is a well-characterized response mechanism against *Toxoplasma* in human fibroblasts^14^. In addition, macrophages from patients with defects in IFNγ-receptor (*IFNR1*) activity have a reduced capacity to kill *Toxoplasma*^15^.

Unlike IFNγ, the role of type I IFNs in human cell-autonomous responses to *Toxoplasma* is still emergent. Type I and type II IFNs activate some overlapping pathways, however, there are type I- and type II-specific ISGs, reflecting the activation of different signalling pathways. In particular, IFNγ induces more genes related to apoptosis and cytokine interactions than IFNα. Additionally, type I IFNs can also inhibit some aspects of IFNγ-induced protective responses^16^. Nevertheless, there is strong empirical evidence that type I IFNs play some role in controlling *Toxoplasma* in human and murine cells. Compared to wildtype controls, mice lacking functional type I IFN receptor (*Ifnar1*) exhibit increased susceptibility to *Toxoplasma*^17^. In addition IFNβ inhibits *Toxoplasma* in human macrophages^18^, retinal epithelial cells^19^, and mouse macrophages^20^. Although these findings suggest a role for type I IFNs in controlling *Toxoplasma*, a systematic analysis of type IFN-induced ISGs with anti-*Toxoplasma* properties in human cells is largely still lacking.

Since IFNs directly and indirectly induce the expression of several genes, the regulatory role of IFN in *Toxoplasma* infection is expected to depend on the expression of ISGs. Recently, genome-wide CRISPR/Cas9 knockout (GeCKO)^21^ or ISG-overexpression^22,23^ screens have been used to rapidly characterize murine or human ISGs. However, the ISG-expression library has limitations in scope and precision. The limitations in scope arise from the limited number of ISGs in the expression library (∼600), which is less than the number of known ISGs in the human genome. Limitations in precision arise from the potential for positive feedback via IFN signalling triggered by overexpression of individual ISGs, which can complicate the ability to precisely identify ISGs that directly control *Toxoplasma*. Similarly, because of the potential effect of genes that function downstream of ISGs, the GeCKO screens cannot distinguish direct and indirect effects of ISGs on *Toxoplasma* growth. Additionally, by targeting all protein coding genes in the human genome, the data analysis in GeCKO screens suffers from a potentially insurmountable multiple-testing problem. Recently a CRISPR/Cas9-based knockout library that targets only type I-inducible ISGs in the human genome (1,902 genes in total) was developed to overcome these limitations and used to systematically identify key components of IFNα-induced antiviral factors in human cells^24^. The library relies on the use of lentiviral vectors to deliver CRISPR/Cas9 and single guide RNAs (sgRNAs) targeting individual ISGs (ISG-knockout library). While this approach has proven highly successful for identifying ISGs that potently inhibit viruses, similar screening methodologies have not yet been adapted for parasites. Here, we performed a loss-of-function screen using the ISG-knockout to rapidly identify ISGs that control *Toxoplasma* in IFNα-stimulated THP-1 cells. The screen revealed known and previously unappreciated inhibitors of *Toxoplasma* including *GBP5, MAX* and *SXN5*. Taken together, these findings reveal effector molecules involved in the complex relationship between *Toxoplasma* and the IFNα response pathway, and open new avenues for exploring the IFNα-induced cell-autonomous immune regulation of *Toxoplasma*.

## RESULTS AND DISCUSSION

### Fluorescence-based loss-of-function screening approach

We employed a loss-of-function screen to identify type I interferon-stimulated genes (ISGs) that control *Toxoplasma* in human macrophages. First, we optimized the screening conditions by determining the suitability of a cytometry-based protocol to quantify infected cells as previously described^25^. To do this, GFP-expressing type I (RH strain) *Toxoplasma* parasites were added to THP-1 cells at a multiplicity of infection (MOI) of 1 and allowed to infect the cells for 1 hour before rinsing out extracellular parasites. Cells were then cultured in fresh cell medium for 2, 12, and 24 hours, providing a temporal evaluation of parasite growth. RH is a type I strain that is susceptible to IFNs in human cells^26^ but compared to other clonal *Toxoplasma* strains replicates faster, thus making it ideal for identifying ISGs that restrict *Toxoplasma* growth. As shown in **Figure 1A**, we observed an increase in the percentage of GFP positive (GFP+) cells over time, indicating that GFP fluorescence can be used to follow parasite growth dynamics in THP-1 cells.

**Figure 1:**
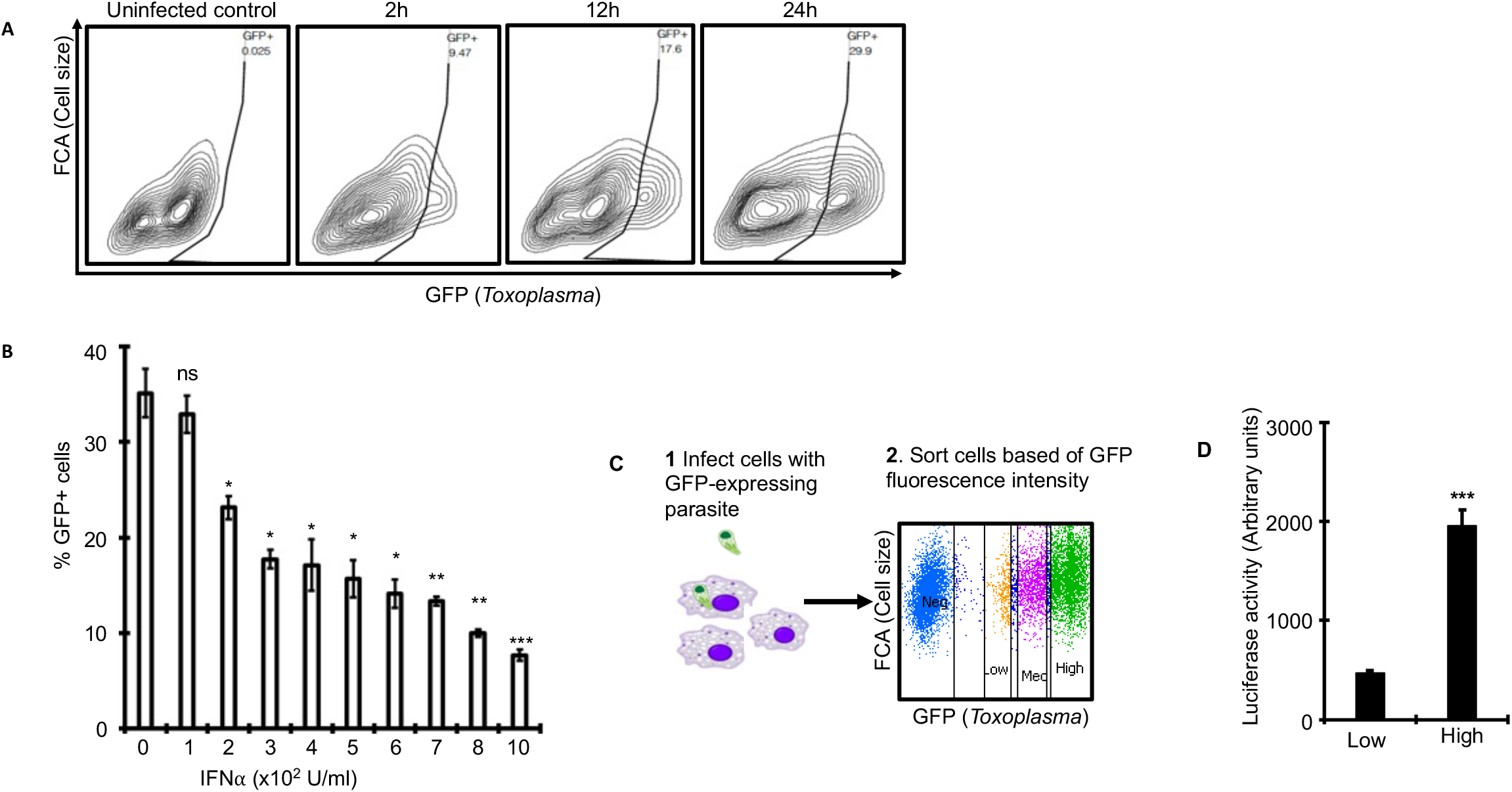
High throughput parasite growth assay. **A**) Representative flow cytometry plots of universal interferon alpha (uIFNα)-stimulated THP-1 cell synchronously infected with GFP-expressing Type I (RH) *Toxoplasma* stains at 2, 12, and 24 h post-infection. Values in the upper right corner of each plot indicate the percentage of GFP-positive cells in singlet cell population. The uninfected control is presented on the left. **B**) Quantification of parasite burden in THP-1 cells stimulated with different concentrations of uIFNα. Data are mean ± s.d. (n = 3, * *p = value* ≤ 0.05, ** = *p value* ≤0.01, ns = not significant relative to control). **C**) THP-1 cells infected with *Toxoplasma* parasites constitutively expressing GFP and firefly luciferase for 24h were sorted in flow cytometry into GFP-negative, GFP-low, GFP-medium, and GFP-high. **D**) Luciferase activity assay, as a proxy of parasite burden, confirm that GFP-high THP-1 cells contain significantly more parasites than GFP-low cells. Data are mean ± s.d. pooled from three independent experiments (*** = *p value* ≤0.001 relative to control).

Next, we sought to define the concentration at which type I interferons minimally inhibit *Toxoplasma* in THP-1 cells. To do this, THP-1 cells pre-stimulated for ∼16h with increasing concentrations of universal IFN alpha (uIFNα), were exposed to syringe-lysed RH parasites for 1 hour before removing extracellular parasites and incubating the cells further in fresh media for approximately 24 hours. Flow cytometry analyses of parasite burden based on GFP fluorescence showed that uIFNα inhibited *Toxoplasma* in a dose-dependent manner, with 200U/ml being the concentration of uIFNα that minimally inhibit the parasite (**Figure 1B**). Finally, we sought to determine if, rather than the overall increase in number of GFP+ cells, we could distinguish THP-1 cells with high and low parasite burden based on GFP fluorescence intensity. As shown in **Figure 1C**, we were able to distinguish cells in the top and bottom 10% of GFP fluorescence intensity spectrum, corresponding to cells with high and low parasite burden, respectively. Finally, because the parasites constitutively express luciferase, we used luciferase activity readout as a proxy of parasite burden, to confirm that the GFP-low and high cells, indeed contained low and high parasite numbers, respectively (**Figure 1D**). Together, these results confirm the suitability of flow cytometry to rapidly measure parasite burden and the inhibitory capacity of uIFNα on *Toxoplasma* in THP-1 cells.

### Unbiased ISG screen identifies regulators of *Toxoplasma* infection

Having established the suitability of flow cytometry to distinguish cells with low and high parasite burden, we next leveraged this approach to identify ISGs that control *Toxoplasma* growth in THP-1 cells. Although several ISGs can inhibit *Toxoplasma*, the parasite can also induce the expression of some ISG to promote its survival and replication in host cells. For example, although interferon regulatory factor 3 (IRF3) is needed by mammalian cells to inhibit many microbes, especially viruses, *Toxoplasma* activates IRF3 to induce the transcription of several genes that in turn promote parasite growth in murine cells^27^. Therefore, we coupled the high throughput cytometry-based parasite growth assay with a CRISPR/cas9-based ISG loss-of-function screen to identify ISGs that control *Toxoplasma* in human THP-1 cells. Briefly, THP-1 cells were transduced with lentivirus expressing CRISPR/Cas9, sgRNA targeting all known type I IFN-inducible ISGs (1,902 genes), 200 non-targeting sgRNAs (controls), and puromycin resistance marker^24^ (Materials and Methods). Although the loss-of-function library is designated as type I ISG knockout library, it is worth noting that some ISGs are induced by both type I IFN and type II (IFNγ) due to the overlap in signaling pathways between type I and type II IFNs. Transduced cells were selected in media supplemented with puromycin for 14 days then stimulated with 200 U/ml of uIFNα, a concentration that minimally inhibit *Toxoplasma* in THP-1 cells, for 24 hours. We used the minimal uIFNα concentration to avoid overriding the effect of uIFNα that would potentially occur at high IFNα concentrations due to feedback via IFN signaling. The cells were then synchronously infected with GFP-expressing RH parasites for 1 hour before removing extracellular parasites and incubating the cells for 24 hours in cell culture media supplemented with dextran sulfate to inhibit late or re-infection by egressed parasites^28^. Next, the cells were sorted into two populations; top 10% of cells expressing the highest GFP fluorescence (GFP-high) and bottom 10% of cells expressing low GFP (GFP-Low) reasoning that these would contain sgRNAs targeting inhibitory and non-inhibitory ISGs, respectively, in the stimulated THP-1 cells (**Figure 2A**). We quantified sgRNA abundance within the sorted populations by high throughput sequencing and detected >95% of sgRNAs in the library, which were relatively uniformly distributed across replicate samples, indicating a constant sgRNA coverage across the different experimental steps. Next, we calculated the abundance-based rank difference between the inhibitory (GFP-High) and non-inhibitory (GFP-Low) fractions to identify sgRNAs that control *Toxoplasma* in uIFNα-stimulated THP-1 cells. We found that sgRNAs targeting guanylate binding protein (*GBP*) 5, a known inhibitor of *Toxoplasma* in THP-1 cells^29^, was enriched in the GFP-high cell population, indicating that the screen can identify regulators of *Toxoplasma* in human cells. In total, we found that sgRNAs targeting 26 ISGs, were significantly (log2FC ≥ 1 and *FDR < 0*.*1*) enriched in the GFP-high, relative to GFP-low, cell population (**Figure 2B**). Meanwhile sgRNA targeting *IRF2*, which can potentially function as a transcriptional inhibitor or activator of the inflammatory IRF1 in different inflammatory states^30^, was enriched in the GFP-low fraction (**Figure 2B**). *Toxoplasma* can also dysregulate ISGs in naïve host cells to promote its survival and growth in host cells^18,31,32^. We reasoned that *Toxoplasma* should be able to replicate robustly in naïve cells, except in cells lacking a parasite-induced gene required to support parasite growth. Therefore, to identify *Toxoplasma*-induced ISGs that potentially promote *Toxoplasma* growth in THP-1 cells, we infected naïve THP-1 cells then sorted 10% of cells expressing the highest and lowest GFP fluorescence. sgRNAs targeting 32 ISGs were significantly (log2FC ≥ 1 and *FDR < 0*.*1*) enriched in the GFP-high relative to GFP-low fraction. These included sgRNAs targeting adenosine monophosphate deaminase (*AMDP3*), 5’-aminolevulinate synthase 1 (*ALAS1*), and phosphoribosylglycinamide formyltransferase (*GART*) enzymes. At least 5 out of 8 sgRNAs targeting these genes had similar effects, suggesting that the phenotypes were not due to off target effects.

**Figure 2:**
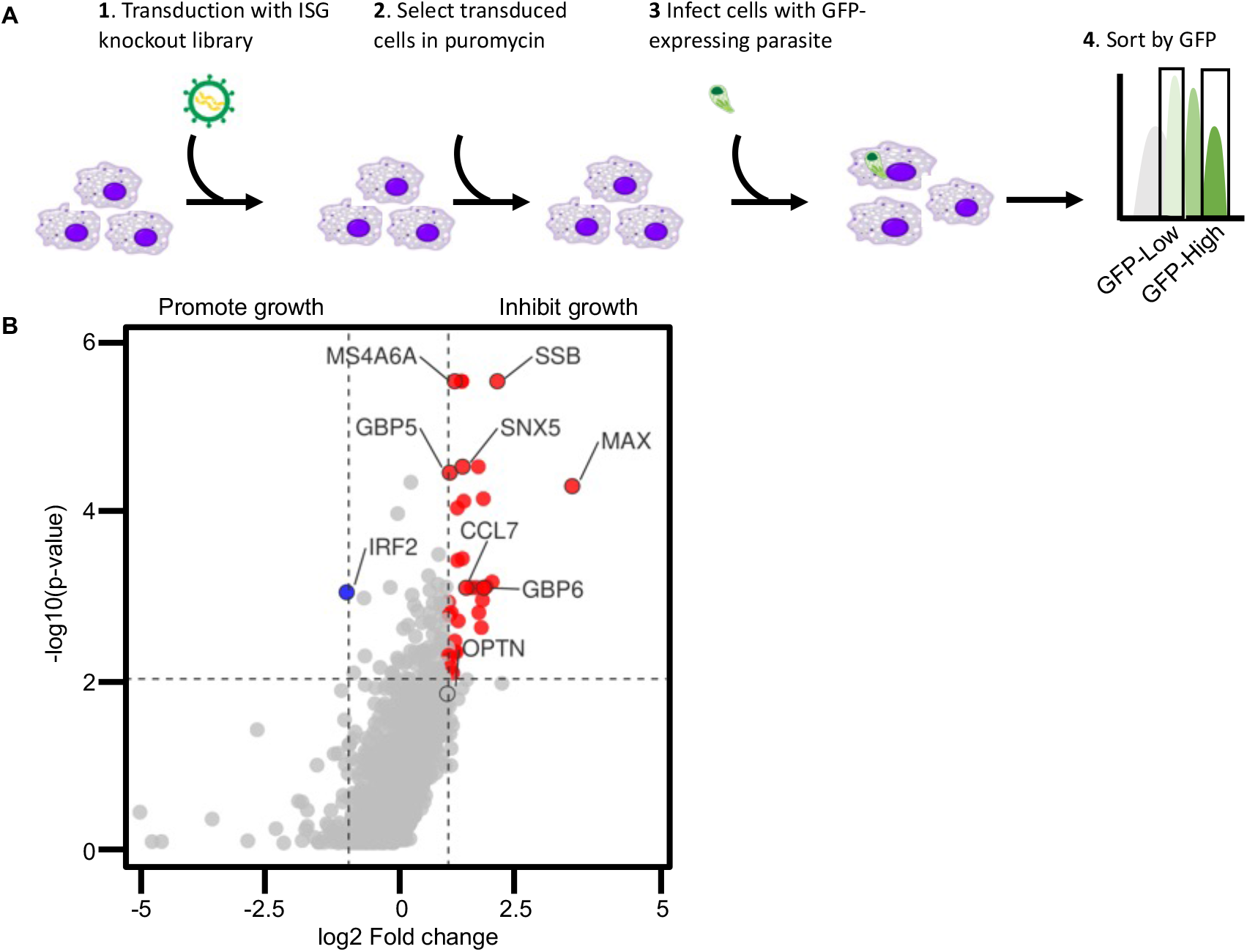
A CRISPR screen for regulators of IFNα-dependent control of *Toxoplasma.* **A**) THP-1 cells transduced with Human Interferon-Stimulated Gene CRISPR Knockout Library was stimulated with uIFNα (200U/mL) for 24h and infected with GFP-expressing type I (RH) *Toxoplasma* parasites. 24 h post-infection, the bottom (GFP-low) and top (GFP-high) GFP-expressing cells were separately collected in a flow cytometer. Genomic DNA was extracted from the sorted cell fractions, sequenced and sgRNA enrichment in GFP-high relative to GFP-low cell fractions determined. **B**) Volcano plot of genes enriched in the top 10%, relative to bottom 10% of GFP-expressing cells. Significantly enriched genes (log2FC ≤1 and FDR<0.1) are shown in red. Depleted genes are shown in blue.

To gain further insights into the hits from the screen, we performed a functional enrichment analysis on the ISGs targeted by the significantly enriched sgRNAs. We found that ISGs that inhibit *Toxoplasma* in stimulated THP-1 cells were enriched in molecular pathways associated with protein heteromerization (*Adj p value* = *0*.*009*), positive regulation of IL1β production (*Adj p value* = *0*.*04*), and SUMOylation of transcription factors (*Adj p value* = *0*.*03*). Functional enrichment analysis of ISGs enriched within the GFP-Low fraction in the non-stimulated cells revealed significant overrepresentation of genes in the purine and heme metabolism pathway, suggestive of important roles for these pathways in *Toxoplasma* growth in THP-1 cells. Taken together, these results demonstrate that the ISG-knockout screen can be used to rapidly identify ISGs that control *Toxoplasma* growth in human cells.

### Validation of inhibitory factors

To validate some of the ISGs that control *Toxoplasma* in THP-cells, we measured parasite burden in THP-1 cells expressing sgRNAs targeting each candidate ISG in a one gene at a time format. *GBP5* is a well-characterized negative regulator of *Toxoplasma* that is reported to reduce parasites burden in THP-1 cells^33^. Compared to wildtype cells, ablation of *GBP5*, as previously described^33^, diminished the ability of uIFNα-stimulated THP-1 cells to inhibit *Toxoplasma* (**Figure 3**). To validate previously unrecognized regulators of *Toxoplasma* in THP-1 cells from the ISG screen, we measured parasite burden in uIFNα-stimulated THP-1 cells that express sgRNA targeting either Small RNA binding exonuclease protection factor La (*SSB*), MYC associated factor X (*MAX*), Sorting nexin 5 (*SNX5*), or Coagulation factor II thrombin receptor like 2 (*F2RL2*), which were ranked in the top 10 of ISGs with significant inhibitory effect on *Toxoplasma* in uIFNα-stimulated THP-1 cells. Compared to cells transduced with control lentivirus, editing *MAX, SNX5, F2RL2*, and *SSB* abolished the inhibitory effect of uIFNα on *Toxoplasma* growth in THP-1 cells (**Figure 3**). *SNX5* binds phosphoinositides such as phosphatidylinositol 3-phosphate (PtdIns(3)P) and is essential for sensing and driving membrane curvature^34^. PtdIns(3)P generation by class III phosphatidylinositol 3-kinase (PI3KC3) complex and membrane remodelling are critical early steps during autophagosome biogenesis^35^. Since autophagy is known to be a key mechanism by which human cells control *Toxoplasma* infections^36,37^, it is possible that *SNX5* control *Toxoplasma* by augmenting autophagy in THP-1 cells. Canonical autophagy involves the formation of double membrane phagophore and the processing and conjugation of the cytosolic associated protein light chain 3 (LC3) to the phosphatidylethanolamine (PE)^36^. Therefore, prior to confirming the impact of *SNX5* on autophagy as a mechanism for regulating *Toxoplasma* in uIFNα-stimulated THP-1 cells, we will need to directly investigate the localization of LC3 on the PVM in *SNX5*-deficient cells relative to wild type cells. The recruitment of LC3 to the PVM in HeLa was shown to be parasite strain-specific, with the type II strain being more susceptible to LC3 coating than the type I used in this study^36^. Human anti-*Toxoplasma* cellular mechanisms are known to be cell-type specific, thus, it is possible that in THP-1 cells, both the type I and II parasite strains are susceptible to autophagy-mediated restriction. *F2RL2* encodes a member of the protease activated receptor (PAR) family, PAR3, which are a class of G protein-coupled transmembrane receptors activated via the proteolytic cleavage of their N-termini and are involved in a number of inflammatory and infectious diseases^38^. Previous studies have shown the involvement of PARs in Toxoplasma-induced intestinal inflammation in mice. Mice lacking a functional PAR2 exhibited reduced intestinal inflammation accompanied by diminished secretion of pro-inflammatory cytokines such as IL6 and CXCL1^39^. It is possible that the anti-*Toxoplasma* activity of *F2RL2* in THP-1 cells is mediated via its ability to regulate uIFNα-induced inflammatory signalling. Like other *Toxoplasma*-restriction factors validated, we do not know whether the effect of *SNX5* or *F2RL2* on intracellular parasite burden is due to their impact of parasite survival or replication. We will also need to determine whether any of the identified restriction factors localize to the PVM or interfere with the localization of other anti-*Toxoplasma* effectors, such as GBPs, to the PVM.

**Figure 3:**
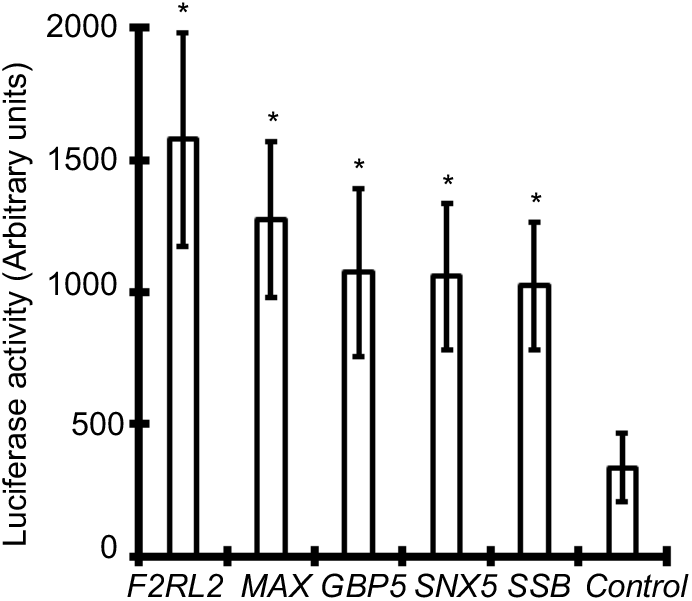
Validation of CRISPR screen hits. Average luciferase activity (a proxy for parasite burden) in uIFNα-stimulated cells expressing sgRNA targeting the indicated candidate ISG. Data are mean ± s.d pooled from three independent experiments relative to uIFNα-stimulated cells expressing sgRNA targeting *ACTB* (* = *p value* ≤0.05 determined by *Student t test*).

## CONCLUDING REMARKS

Type I interferon (IFN) is a multi-gene cytokine family that encodes thirteen partly homologous IFNα subtypes in humans, one IFNβ, and several poorly defined single gene products^4^. This work focused on IFNα, which together with IFNβ, are well characterized and broadly expressed. Compared to type II interferons (IFNs), the role of type I IFNs in human cell-autonomous responses to *Toxoplasma* is not well established. To address this gap in the knowledge, we adapted a flow cytometry-based loss-of-function screening approach to identify cellular regulators of *Toxoplasma* infection among 1902 type I IFN-stimulated genes (ISGs) in THP-1 cells. We elected to use parasite fluorescent intensity as our screening output as it is high throughput and a broad phenotype governed both by parasite death and growth inhibition. The screen identified both known (for example Guanylate binding protein 5, *GBP5*) and previously unappreciated cell-autonomous inhibitors of *Toxoplasma*. The findings were confirmed by knocking out five candidate ISGs in THP-1 cells and we are continuing to validate and functionally characterize more candidate ISGs. This report provides a genomic landscape of the regulation of IFNα pathway in THP-1 cells and its impact on *Toxoplasma*, and potentially other intracellular pathogens.

## MATERIALS AND METHODS

### Parasites and Cell culture

The type I (RH) *Toxoplasma* parasites engineered to express green fluorescent protein (GFP) and firefly luciferase have previously been described^40^. The parasites were maintained by serial passage on confluent human foreskin fibroblast (HFF) monolayer. THP-1 cells (TIB-202), purchased from ATCC, were cultured in Roswell Park Memorial Institute (RPMI) media (Invitrogen) supplemented with 10% heat-inactivated foetal bovine serum (FBS) (Invitrogen), 10 mM HEPES buffer, pH 7.5, 2 mM L-glutamine, and 50 μg/ml penicillin (Thermo Fisher Scientific). Cells were incubated at 37°C in 5% CO_2_.

### Lentivirus production

Lentivirus stocks expressing the Human ISG CRISPR Knockout (ISG-knockout) library components were produced as previously described^24,41^, with slight modifications. Briefly, HEK293 T cells (ATCC) were seeded at 18 million cells in T175 flasks the day before transfection in Dulbecco’s Modified Eagle Medium (DMEM) supplemented with 10% FBS. One hour prior to transfection, media was removed and replaced with pre-warmed OptiMEM media (Life Technologies). Cells were transfected with 20μg of ISG-knockout llibrary containing 15416 sgRNAs (Addgene, Cat # 125753), 10μg and 15μg of second-generation lentiviral packaging plasmid psPAX2 (Addgene, Cat # 12260) and pMD2.G (Addgene, Cat # 12259), respectively, using Lipofectamine 3000 and Plus reagent (Life Technologies) according to the manufacturer’s protocol. Six hours after transfection, the OptiMEM media was replaced with 30 ml DMEM supplemented with 10% FBS. The following day, the media was replaced with fresh media supplemented with 500x viral boost reagent (Alstem) as per the manufacturer’s recommendations. At 48 hours post transfection, the viral supernatant was collected and centrifuged at 300 g for 10 minutes and passed through a 0.45 μm filter to remove cell debris. The lentiviral particles were then concentrated by adding virus precipitation solution (Alstem) to the viral supernatant followed by incubation overnight at 4^°^C and centrifugation at 1500 g for 30 minutes at 4^°^C. The lentiviral pellet was resuspended at 100x of original volume in cold DMEM and stored at -80^°^C until use.

### Lentiviral transduction and screening

THP-1 cells were transduced in triplicates using the TransDux MAX transfection reagent (System Biosciences, LV860A-1) according to the manufacturer’s recommendations. Briefly, 5 million THP-1 cells were seeded in 10cm petri dishes in RPMI supplemented with TransDux MAX transfection reagent. Lentivirus was then added to the cells at a 1:250 v/v ratio and incubated for 24h at 37 °C, 5% CO_2_. The media was then removed, and the cells cultured in RPMI media supplemented with puromycin for 14 days with frequent passage to select for transduced cells and generate enough cells for the screening experiments.

Three million cells from each triplicate were seeded separately in 6 well-plates in fresh RPMI media and left unstimulated or stimulated with universal interferon alpha (200U/ml) (R&D systems) for 24 hours. The cells were then left uninfected or infected with syringe-lysed GFP-expressing RH parasites at a multiplicity of infection (MOI) of 1, using a temperature shift-based synchronized invasion approach^42^. Briefly, *Toxoplasma* parasites were syringe-lysed using 25G needles, passed through 5μM filter to remove HFFs, and centrifuged at 500 x g for 7 minutes. The parasites were then resuspended in fresh RPMI medium, added to the THP-1 cells at MOI of 1 room temperature, and immediately centrifuged at 250 g for 3 minutes followed by incubation at room temperature for 5 minutes. The cells were then transferred into 37 °C, 5% CO_2_ incubator. Two hours post-infection, extracellular parasites were removed with three phosphate buffered saline (PBS) washes before culturing the cell further in RPMI supplemented with 10% FBS and Dextran sulfate, to inhibit re-infection or late infections^28^. At 24 hours post infection the cells were harvested and processed for flow cytometry.

### Flow cytometry

Three million infected or uninfected THP-1 cells were harvested and washed twice with warm PBS. Cells were resuspended in PBS + 1% bovine serum albumin (BSA). All samples were analyzed on a LSR Fortessa (BD Biosciences), and recorded data was processed using FlowJo 10.3 (FlowJo, LLC). To perform the screen, 500,000 representative cells of GFP-Low (bottom 10%), and GFP-High (Top 10%), were sorted into lysis buffer for genomic DNA extraction. At the same time 500,000 viable uninfected cells were sorted into lysis buffer using the LIVE/DEAD dye. 500,000 unsorted cells from each population (infected and uninfected) were used as input controls.

### Preparation of genomic DNA for sequencing

Genomic DNA (gDNA) was isolated separately from the sorted and unsorted cell fractions using DNeasy Blood and Tissue Kit (Qiagen). Amplification and bar-coding of sgRNAs was performed as described previously^24,41^. Briefly, each sample was divided into 10 50μl reactions with 2μg gDNA, 2x High Fidelity PCR Master Mix, 10μM of both forward and reverse primer 1. The PCR primers and conditions are available from Addgene. The PCR product were mixed and cleaned using QIAquick PCR Purification Kit (Qiagen). Next, 5μl was taken from the cleaned PCR product to go into a second PCR for indexing. The indexed PCR products were cleaned using 1 Volume AMPure XP beads on magnetic stand and eluted with EB buffer (QIAGEN) followed by quantification using a Qubit dsDNA HS Assay Kit. The product size was also confirmed by running 2ul of the PCR product on a 2% agarose gel. Purified samples were multiplexed and sequenced on a NextSeq 500 machine to generate ∼40 million 75-bp single-end reads per sample (Edinburgh Clinical Research Facility).

### Analysis of ISG CRISPR Screens

Sequencing reads were processed and analyzed using the MAGeCK software^43^ to identify negative and positive hits in the screens by quantifying and testing for sgRNA enrichments as previously described^24,41^. Briefly, sgRNA abundance was first determined using the MAGeCK ‘count’ module for the raw sequencing reads using default settings and removing sgRNAs with less that 50 reads in two out of the three replicates. The MAGeCK ‘test’ module, was then used to normalize the samples for varying sequencing depths and to test for sgRNA and gene-level enrichment. The non-targeting sgRNAs controls in the ISG-knockout library were used to estimate size factor for normalization and build the mean variance. The sgRNA-level enrichment scores from MAGeCK were then used to generate gene-level enrichment scores by alpharobust rank aggregation (RRA). Significantly positively or negatively selected ISGs were selected based on absolute log2 fold change ≥ 1 and Bonferroni-corrected false discovery rate (FDR) < 0.1.

### Gene Set Enrichment Analysis for Screen Hits

We used Gene Set Enrichment Analysis (GSEA) in the fgsea package^44^ to identify enriched annotations in the ISGs that were significantly positively or negatively selected for in the screens using logFC values for all ISGs in tested in the screen. We used the KEGG pathways dataset as the reference gene annotation database. Normalized enrichment scores and p-values were determined by a permutation test with 10,000 iterations with same size randomized gene sets and adjusted with the FDR method.

### Functional validation candidate genes

The sequences of gRNAs used for CRISPR knockout for *GBP2* and *SNX5* have previously been described^29,45^. The corresponding top ranked sgRNAs from the screen were used to individually knockout *MAX, SSB*, and *F2RL2*. For generation of knockout cell lines, sgRNA targeting each gene was individually cloned into the pLentiCRISPR-V2 backbone^46^ and transfected into 293T cells along with packaging plasmids (psPAX2 and pMD2.G) to generate sgRNA- and Cas9-expressing lentivirus. THP-1 cells were separately transduced with lentivirus expressing sgRNA targeting each candidate gene and selected with 5 μg mL-1 Puromycin (A1113802, Gibco) for 7 days; all control (non-transduced cells in Puromycin) were dead by this time. The transduced cells were sub-cloned by serial dilution into 96-well plates using complete medium supplemented with non-essential amino acids (11140076, Gibco), penicillin/streptomycin and GlutaMAX. The clones were expanded into 6-well plates and the frequency of deletions and insertion in the targeted gene confirmed using deep targeted sequencing, which confirmed the sgRNA-targeted deletion of each gene in THP-1 cells.

## AUTHOR CONTRIBUTION

MH conceived the project. AG, AR, and MH designed and conducted the experiments and wrote the manuscript.

